# Voltage-gated potassium channels ensure action potential shape fidelity in distal axons

**DOI:** 10.1101/2020.09.15.297895

**Authors:** Victoria Gonzalez Sabater, Mark Rigby, Juan Burrone

## Abstract

The initiation and propagation of the action potential (AP) along an axon allows neurons to convey information rapidly and across distant sites. Although AP properties have typically been characterised at the soma and proximal axon, the propagation of APs towards distal axonal domains of mammalian neurons remains limited. We used Genetically Encoded Voltage Indicators (GEVIs) to image APs simultaneously at different locations along the long axons of dissociated hippocampal neurons with sub-millisecond temporal resolution. We found that APs became sharper and showed remarkable fidelity as they traveled towards distal axons, even during a high frequency train. Blocking voltage-gated potassium channels (K_v_) with 4-AP resulted in an increase in AP width in all compartments, which was stronger at distal locations and exacerbated during AP trains. We conclude that the higher levels of Kv channel activity in distal axons serves to sustain AP fidelity, conveying a reliable digital signal to presynaptic boutons.

## Introduction

Projection neurons in the central nervous system extend long axons often forming thousands of small en-passant synaptic boutons, hundreds of microns from their cell soma (Debanne et al., 2011). The probability that neurotransmitter is released from such boutons (Pr) depends on a number of factors, which are set in motion by the arrival of an action potential (AP) (Branco and Staras, 2009; Dittman and Ryan, 2019; Kawaguchi, 2019). Modest changes in AP waveform have the capacity to strongly influence voltage-gated Ca2+ channel open probability (Scarneti et al., 2020), the subsequent driving force for Ca2+ entry (Scarnati et al., 2020), and given its highly non-linear calcium-dependence (Schneggenburger and Neher, 2000; Neher and Sakaba, 2008), the probability of release (Rama et al., 2015). A model of synaptic transmission from distally-located projection neuron release sites therefore requires knowledge of AP propagation and the AP waveform.

The AP is not an immutable waveform from initiation to termination but is appreciated to vary as it propagates along the axon (Branco and Staras, 2009; Debanne et al., 2011; Scarneti et al., 2020). A handful of mostly pipette-based electrophysiological studies have suggested that variations in axon morphology, as well as Nav and Kv channel expression, confer characteristics to the axonal AP waveform, which differ from the soma, and potentially within different regions of the axonal tree (Goldstein and Rall, 1974; Geiger and Jonas, 2000; Kole, Letzkus and Stuart, 2007; Hoppa *et al*., 2014; Cho *et al*., 2017). Such local properties have been shown to locally impact axonal AP propagation fidelity (Khaliq and Raman, 2005; Monsivais, 2005; Sasaki, Matsuki and Ikegaya, 2012; Kawaguchi and Sakaba, 2015; Cho *et al*., 2017), speed (Chéreau et al., 2017), and shape (Kole, Letzkus and Stuart, 2007; Alle, Kubota and Geiger, 2011; Hoppa *et al*., 2014). Moreover, the axon has been shown to exhibit a range of NaV inactivation (Rama *et al*., 2015; Zbili *et al*., 2020), and susceptibility to Kv channel modulation linked to frequency-dependent AP broadening (Geiger and Jonas, 2000) or attenuation (Kawaguchi and Sakaba, 2015), as well as depolarisation-induced analogue facilitation (Kole, Letzkus and Stuart, 2007; Shu *et al*., 2007; Foust *et al*., 2011; Bialowas *et al*., 2015). It is therefore important to understand both the spatial and temporal modulation of APs as they travel along an axon and during high frequency firing regimes.

The localised heterogeneity in axonal AP properties ideally requires the capacity to record the AP at multiple locations. Whilst traditional pipette-based strategies offer unparalleled temporal and voltage sensitivity, they generally allow for only single-site measurements, and their paucity is attributable to the technical difficulty of recording from small structures (typically less than 1 µm) (Novak *et al*., 2013; Kawaguchi and Sakaba, 2017; Vivekananda *et al*., 2017). Furthermore, given whole-cell patch-clamp techniques both dialyses the internal cytosol, and rupture large parts of the membrane under study, alternative higher-throughput approaches are worth consideration.

Optical reporters of membrane voltage offer a potential solution that circumvent some of the pitfalls of patch-clamp recordings. Voltage-sensitive dyes (VSDs) have been previously employed for voltage imaging in sub-cellular compartments (Sabatini and Regehr, 1997; Popovic *et al*., 2011; Rowan, Tranquil and Christie, 2014). Although genetically-encoded voltage indicators (GEVIs) have traditionally lagged behind the VSDs in their performance, recent advances in the development of GEVIs have opened the possibility of recording changes in voltage in sub-cellular compartments non-invasively and with improved temporal resolution (Panzera and Hoppa, 2019). For instance, Hoppa et al. 2014 used a configuration with sufficient sensitivity to report differences in the AP waveform within distally located hippocampal boutons (Hoppa *et al*., 2014).

More recently, GEVIs with greatly improved properties have been developed, but have not been validated beyond their original reports. To image rapid events such as an AP in small sub-cellular compartments, the probe must have the necessary temporal resolution and sufficient brightness to resolve the AP. Although there are many new GEVIs available (Platisa and Pieribone, 2018; Knöpfel and Song, 2019), few of them meet these requirements. One notable group of GEVIs, based on non-pumping variants of microbial rhodopsins, have particularly fast kinetics (in the sub-millisecond range) (Arch; Kralj *et al*., 2012; Hochbaum *et al*., 2014), and exhibit sufficient voltage sensitivity to image the action potential in boutons (Hochbaum *et al*., 2014; Hoppa *et al*., 2014), but are not particularly bright. However, a more recent screen of microbial rhodopsin mutants generated new brighter voltage sensors, including Archon1 and Archon2 (Piatkevich *et al*., 2018). Whilst Archon1 was capable of responses with large signal-to-noise and negligible photo-bleaching, Archon2 was more suited to imaging the action potential in axonal varicosities given its 8-fold increase in brightness compared to Archon1, and fast response kinetics (Piatkevich *et al*., 2018).

Another strategy for generating GEVIs has been to isolate the voltage sensing domain from an opsin, and couple it to a brighter fluorophore through electrochromic FRET. Of the sensors generated using this approach, Ace-2N-4AA-mNeon was particularly promising given its reported sub-millisecond kinetics, the large quantum efficiency of mNeonGreen, and its photo-stability relative to other FRET-opsin based sensors (Gong *et al*., 2015). As a result, and based on their reported brightness and kinetic properties, we selected Ace-2N-4AA-mNeon and Archon2 for a side-by-side comparison of their ability to measure the AP waveform and propagation along an axon. We then utilised the spatial and temporal resolution provided by the GEVIs to characterise differential local properties of the AP as it propagates along the soma and the axonal arbour. We show that AP width sharpens towards distal axonal domains and that the shape of the AP remains largely unaltered during a 20 Hz AP train in distal but not in proximal axons or in the soma. Furthermore, we show that Kv channels in distal axons limit the width and amplitude of an AP thereby improving AP reliability during high frequency trains.

## Methods

### Hippocampal neuronal cultures and transfection

Hippocampi were dissected from embryonic day 17.5 Wistar rat pups of either sex, treated with trypsin (Worthington) at 0.5 mg/mL, and mechanically dissociated using fire polished Pasteur pipettes. Neurons were plated on 18 mm glass coverslips (Thermo Fisher Scientific) pre-treated with 100 µg/mL poly-L-lysine (Sigma) and coated with 10 µg/mL laminin (Life Technologies). Cultures were maintained in Neurobasal medium (Life Technologies) with B27 (1x, Invitrogen) and GlutaMAX (1x, Life Technologies), supplemented with foetal bovine serum (FBS, 2%, Biosera) and Penicillin/Streptomycin (1%, Sigma), at 37°C in a humidified incubator with 5% CO2.

For transfections with Ace2N-mNeon-4AA (Ace-mNeon, Gong *et al*., 2015, distributed by Biolife Inc.), the Effectene transfection reagent (Qiagen) was used. The medium was changed at 3 days *in vitro* (DIV) to culture medium without antibiotic or FBS, and transfections with Effectene were performed at DIV 7 following the manufacturer’s protocol. After transfection neurons were maintained in serum-free media without antibiotics. For transfections with Archon2 (Piatkevich *et al*., 2018, gift from E. Boyden lab), the calcium-phosphate method (Ca-Phos) was used. DIV 3-5 neurons were transfected following an adapted version of a low-toxicity protocol for low density cultures (Jiang and Chen, 2006). The coverslips were returned to their original culture medium with FBS for two more days before changing to serum-free media without antibiotics.

For all experiments 30% of the medium was changed weekly and neurons were imaged 7-14 days after transfection (14-21 DIV).

### Live-cell recording and imaging conditions

Neurons were imaged using an inverted Olympus IX71 epifluorescence microscope with a 60x 1.42 NA oil-immersion objective (Olympus, Fig. 2.1). Coverslips were mounted in a heated chamber (Warner instruments; total volume ∼500 µL) and placed on an IMTP microscope stage (Scientifica). Cells were maintained in external HEPES-buffered saline solution (HBS: 2 mM CaCl_2_, 1.6 mM MgCl_2_, 1.45 mM NaCl, 2.5 mM KCl, 10 mM Glucose, 10 mM HEPES, pH=7.4, Osmolarity=290 mOsm). For experiments at physiological temperature the chamber was heated to 32-35°C, and the pH of the HBS solution was adjusted for this temperature.

Current-clamp recordings were made in the whole-cell configuration from the soma of visually-identified transfected neurons at room temperature. Recordings were performed with borosilicate glass pipettes pulled to a resistance of 4-6 MΩ, fire-polished and filled with internal solution (125 mM KMeSO_4_, 5 mM MgCl_2_, 10 mM EGTA, 10 mM HEPES, 0.5 mM NaGTP 5 mM Na_2_ATP, pH=7.4). Data were acquired with a MultiClamp 700B amplifier (Molecular Devices), and digitized with a Digidata 1440A digitizer (Molecular Devices) at a sampling rate of 20kHz. Recordings were acquired using Clampex 10.3 (Molecular Devices) with a gain value of 5 and a Bessel filter set to 10 kHz. Pipette capacitance neutralisation and bridge balance were applied.

For Ace-mNeon imaging experiments, ∼505 nm excitation illumination was provided using a 525nm LED (Solis), a 500/20 nm excitation filter, and a 510 nm long-pass dichroic mirror. Ace-mNeon emission transmitted through the dichroic was filtered using a 520 nm long-pass filters (Chroma). A power density of 10 mW/mm^2^ was obtained at the specimen plane. Excitation of Archon2 was achieved using a 635 nm diode laser (MRL-III-635L >200 mW <1% RMS, ReadyLasers), expanded 3-fold using a custom Galilean beam expander, and focused onto the back aperture of the objective. Excitation illumination passed through a 640/30 nm excitation filter, and a 660 nm long-pass dichroic, emitted fluorescence was filtered using using a 690/50 nm filter (Chroma). The power density achieved for Archon2 imaging was 2.6 W/mm^2^ at the specimen plane.

Images were acquired at 3.2 kHz with an ORCA-Flash4.0 V2 C11440-22CU scientific CMOS camera (Hamamatsu) cooled to ∼-20 °C with the Exos2 water cooling system (Koolance). Images were acquired with HCImage software (Hamamatsu), binned to 4×4 and cropped to a 16 x 512 pixel region of interest, necessary to achieve the high image acquisition rates. Images were saved in CXD format. For the reconstruction of the axonal arbour after the experiment, high resolution 2048 x 2048 images were acquired with an exposure of 100 ms.

Unless paired with whole-cell patch-clamp recordings where AP stimulation was achieved through the patch pipette, stimulation within imaging experiments was achieved with an extra-cellular tungsten parallel bipolar electrode (FHC) mounted on a PatchStar motorised micro-manipulator (Scientifica). For experiments with axonal imaging only, 1 ms pulses of 10 mV were delivered, whilst 50 µs pulses of 30 mV were applied for experiments involving both somatic, and axonal imaging. Shorter pulses were required when imaging the somatic AP to avoid the stimulation pulse artefact from contaminating the AP signal.

The timing of the LEDs and laser, acquisition and stimulation were triggered externally through Clampex software (pClamp 10, Molecular Devices).

Unless otherwise stated, all recordings were performed in presence of NBQX (10 µM, Tocris), Gabazine (10 µM, Tocris) and D-2-amino-5-phosphonovalerate (APV, 25 µM, Tocris) in order to block synaptic transmission and ensure that the observed events were due to the stimulation only. When necessary, 4-aminopyrimidine (4-ap, 30 µM, Tocris) and/or Tetrodotoxin (TTX, 40nM, Tocris) were supplied using a custom gravity-fed perfusion system powered, where bath volume was maintained by a peristaltic pump (Watson-Marlow 120s).

### Electrophysiology and GEVI recording analysis

Analysis of patch-clamp recordings was performed with custom software written in MATLAB (Mathworks). Values of cell capacitance, input resistance, and series resistance were estimated from membrane test recordings performed at regular intervals. All cells with a series resistance >30MΩ were discarded, or ceased to be recorded from. Current-clamp recordings of APs in the soma were aligned to the AP peak, averaged, and the APs were analysed to extract values for amplitude, onset, full width at half maximum (width), 20-80% rise time and 80-20% decay time, measured relative to baseline membrane potential.

Voltage imaging recordings were analysed in ImageJ and custom scripts within MATLAB software as follows. Prior to analysis, if drift had occurred in the x and y axis during the recording, the images were aligned using the TurboReg ImageJ plugin (Thévenaz, Ruttimann and Unser, 1998). Images were then imported into MATLAB and fluorescence intensity profiles were obtained from regions of interest (ROIs) manually drawn around neuronal structures identified visually on a maximum projection image of the time series by averaging across the ROI pixels in each frame. The fluorescence profile was separated into single trials and corrected for background camera noise by subtracting the average intensity value of the timepoints where no illumination light was applied. Then, recordings were reconstructed at a sampling rate of 100 kHz using cubic spline interpolation. Bleaching was estimated by fitting a single exponential function to each individual trace smoothed with an averaging filter with a window of 3 and interpolated. The resulting curve was used to correct an unfiltered interpolated version of the recording. An accurate representation of the AP waveform with an enhanced SNR was obtained by averaging over the repeats and extracting the AP parameters from the resulting trace. Non-aligned averages were used for analysis except for dual patch-clamp and voltage imaging experiments, where the peak of the electrophysiological recording could be used as reference for alignment. The parameters of the AP waveform measured were amplitude (ΔF/F), SNR, width, 20% to 80% rise time, 80% to 20% decay time and decay time constant. The fluorescence profile during the 20 ms preceding a response was used as baseline for the calculations. The SNR was calculated as the ratio of the response amplitude to the standard deviation of the baseline. For some experiments, the coefficient of variation (CV) was calculated as the ratio of the standard deviation of the variable to its mean. The bleaching rate was estimated from the raw fluorescence time profile excluding the timepoints with no illumination.

The axonal arbour reconstruction was done in ImageJ. Overlapping images were stitched together using the MosaicJ plugin, and the axon was then traced with the NeuronJ plugin.

### Statistical analysis

Statistical analysis was performed using GraphPad Prism 8. All data are presented as the mean ± the standard error of the mean (SEM). Datasets were not assumed to be normally distributed unless the data passed a D’Agostino normality test. Results were considered significant at p < 0.05.

## Results

### Measuring the AP waveform with GEVIs

To accurately perform GEVI-based measurements of the AP along the axons of dissociated hippocampal CA1 pyramidal neurons, we first assessed the ability of the selected GEVIs, Ace-2N-4AA-mNeon and Archon2, to reliably report AP waveform at the soma. Somatic APs were evoked by current injection using a patch-pipette, and the resulting APs were recorded in the whole-cell current-clamp configuration. At the same time, the fast changes in GEVI fluorescence expected from an AP signal were imaged at a high frame rate (3.2 KHz) in a subsection of the somatic membrane (Fig. 1A). For both indicators, APs imaged at the soma could be reliably detected in single trials (Fig. 1B). When averaged across 20 trials, the imaged AP traces for both GEVIs closely resembled the electrophysiologically-recorded AP waveform, even reporting small changes in voltage such as the slow, sub-threshold voltage rise preceding the AP (Fig. 1 B-C’’). In general, we found that Archon2 displayed a larger change in fluorescence to single APs than Ace-mNeon (%ΔF/F: 10% and 14% for Ace-mNeon and Archon2, respectively; p<0.001 t test, Supp. Fig. 1 A), while Ace-mNeon showed a higher signal-to-noise-ratio (SNR: 88.9±16 and 43.8±3.7 for Ace-mNeon and Archon2, respectively; p<0.05 Mann-Whitney test, Supp. Fig. 1 B). Notably, Ace-mNeon displayed a small residual photocurrent in response to illumination (505 nm LED at 10 mW/mm^2^), characterised by an initial depolarising transient in membrane voltage (Vm) that stabilised to a smaller depolarising steady-state Vm offset within a few hundred milliseconds (1.51 mV over the resting membrane potential; CI = 0.5 to 2.7; p<0.001 one-sample Wilcoxon test; Supp. Fig. 1 C and D). To avoid the transient change in Vm from contaminating the optical AP signal we proceeded to elicit somatic APs with current injections delivered 500 ms after illumination onset, when a steady-state depolarisation had been reached. No changes in conductance were observed for Archon2-expressing neurons upon illumination (635nm laser at 2.6W/mm^2^; Supp. Fig. 1 C and E; one-sample Wilcoxon tests).

**Figure 1.**
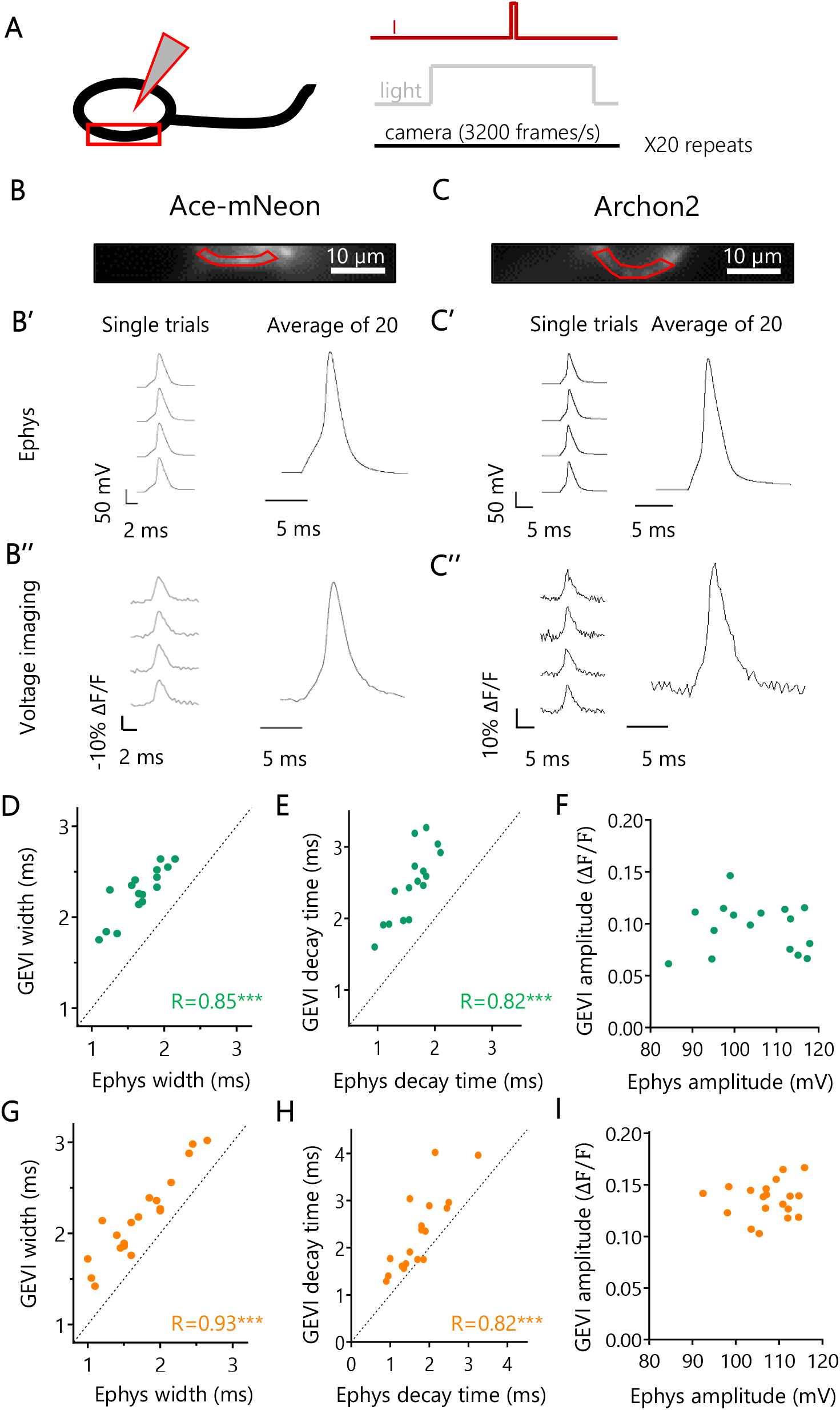
Optical and electrophysiological measure of the somatic AP waveform. **(A)** Schematic representation of the experimental set-up: neurons expressing a GEVI were held in whole-cell current-clamp while simultaneously imaging a segment of their somatic membrane at room temperature. The stimulation protocol consisted in a time-locked current pulse delivered through the patch pipette (I) to evoke a single AP, while subjecting the cells to LED or laser illumination (light) and high-speed camera acquisition at 3.2kHz. The protocol was repeated 20 times and the resulting recordings were averaged. **(B and C)** Representative example cells expressing Ace-mNeon (B) and Archon2 (C), with selected ROIs of the somatic membrane for optical trace analysis shown in red. **(B’ and C’)** Reference Ephys recordings of evoked somatic APs acquired from the example cells in B and C, respectively. Single trials and average traces over groups of 20 trials are shown. **(B’’ and C’’)** Voltage imaging recordings of the somatic AP acquired with Ace-mNeon (B’’) and Archon 2 (C’’), extracted from the ROIs shown in B and C and corresponding to the Ephys recordings in B’ and C’, respectively. **(D-F)** Evoked AP width (D), decay time (E) and amplitude (F) were measured simultaneously with Ace-mNeon and reference Ephys recordings in the same cells to assess accuracy of GEVI measurements. **(G-I)** Evoked AP width (G), decay time (H) and amplitude (I) were measured simultaneously with Archon2 and reference Ephys recordings in the same cells to assess accuracy of GEVI measurements. Ace-mNeon: N=16 cells; Archon2: N=19 cells. R, Pearson’s correlation coefficient; ***, p<0.001. All measurements performed on averages of 20 repeats.

We next examined how the main features of the AP waveform compared across different cells when measured with GEVIs and electrophysiology. For both Ace-mNeon and Archon2, we found that optically-recorded measures of AP width and decay time were correlated with the current-clamp recorded AP, albeit with a small overestimation of the apparent AP kinetics (Fig. 1 D, E, G, H). The overestimation was larger for Ace than for Archon, most likely due to the faster reported kinetics of Archon2 (Supp. Fig. 2 A and B; Piatkevich *et al*., 2018; Gong *et al*., 2015). The AP amplitude showed no correlation between GEVI, and whole cell current-clamp recordings, suggesting that optically-acquired amplitude values cannot be compared across cells (Fig. 1 F and I). A likely explanation for this result is the possible heterogeneity in baseline fluorescence levels resulting from GEVI molecules present within inner membranes, located too far from the plasma membrane to respond to changes in membrane voltage.

Having established the limitations that arise when comparing the AP waveform across different cells, we went on to explore the modulation of AP shape within the same neuron. By performing simultaneous GEVI imaging and current-clamp recordings before and after the addition of voltage-gated channel antagonists (30 µM of the Kv blocker 4-ap and/or 40 nM of the Nav blocker TTX) we were able to subtly alter the AP waveform (Fig. 2). Application of these antagonists either together or separately created a large palette of AP waveforms that we then compared to their corresponding electrophysiological measures. We found that both Ace-mNeon and Archon2 could faithfully report changes in AP kinetics (Fig. 2 C, D, F, G), with relatively small measurement errors (Supp. Fig. 2 C and D). More importantly, we now observed a strong correlation between AP amplitude measured optically and electrophysiologically (Fig. 2 E and H). Together, our data provides strong evidence that Ace-mNeon and Archon2 can be used to measure AP kinetics reliably both within and across cells. AP amplitude, on the other hand, can only be compared within the same membrane segment.

**Figure 2.**
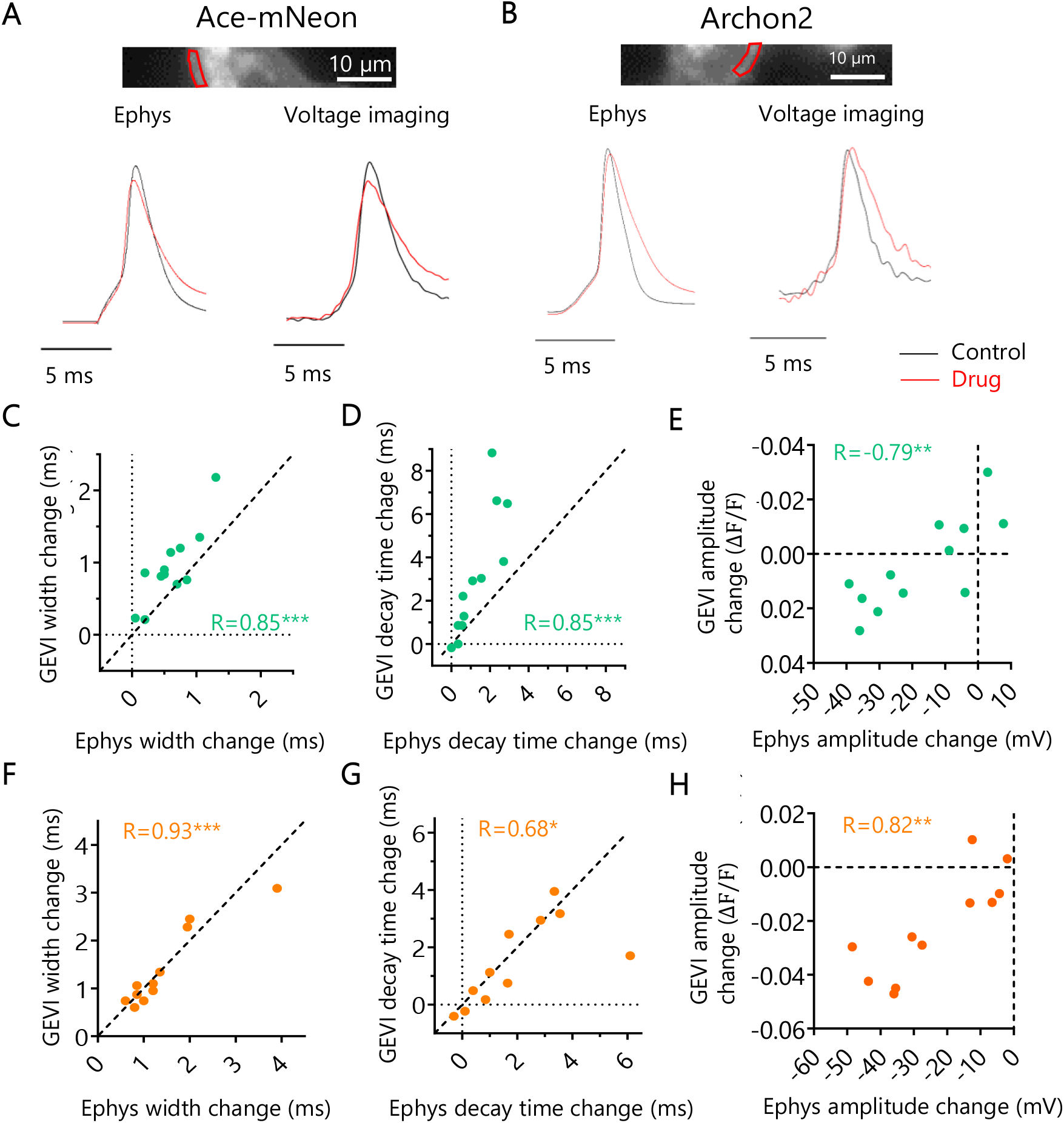
Optical and electrophysiological measure of drug-induced modulation of somatic AP waveform. **(A and B)** Above, imaging window with selected ROI of the somatic membrane shown in red. Below, average traces before and after addition of 30 µM 4-ap and 40 nM TTX (drug), recorded with Ephys and Ace-mNeon / Archon2. **(C-E)** Change in AP width (C), decay time (D) and amplitude (E) induced by perfusion with drug, measured simultaneously with Ace-mNeon and Ephys in the same cells for comparison. **(F-H)** Change in AP width (C), decay time (D) and amplitude (E) induced by perfusion with drug, measured simultaneously with Archon2 and Ephys in the same cells for comparison. Ace-mNeon: N=12 cells; Archon2: N=11 cells. Change calculated as difference between drug and control. R, Pearson’s correlation coefficient; ***, p<0.001; **, p<0.01; *, p<0.05.

Reassured that both GEVIs were able to report changes in AP waveform at the soma with comparable accuracies, we went on to test whether it was possible to image the AP in the axons of dissociated CA1 pyramidal neurons. Compared to the soma, the plasma membrane area in the axon is much reduced, meaning far less GEVI molecules are available to be trafficked there and imaged. The result is a relatively lower baseline fluorescence level from background, which at high acquisition rates meant the GEVI signal was more vulnerable to camera noise (Popovic *et al*., 2015). At near physiological temperatures (32°C) APs were elicited by somatic stimulation with a bipolar electrode and both Ace-mNeon and Archon2-mediated responses were detected in single trials (Fig. 3 A). To enhance the SNR we averaged across multiple repeats (from 20 to 50), with the number of repeats tailored to the levels of photobleaching for each probe (Fig. 3 B and C). When analysing the baseline GEVI fluorescence over time, Ace-mNeon exhibited higher bleaching rates than Archon2, with time constants of 45.71s and 143.62 s, respectively (Supp. Fig. 3 G). Despite the loss of fluorescence over time, the recorded AP width, and amplitude for both GEVIs remained constant for extended periods, allowing for several sets of repeats under different conditions to be performed in a single experiment (Supp. Fig. 4).

**Figure 3.**
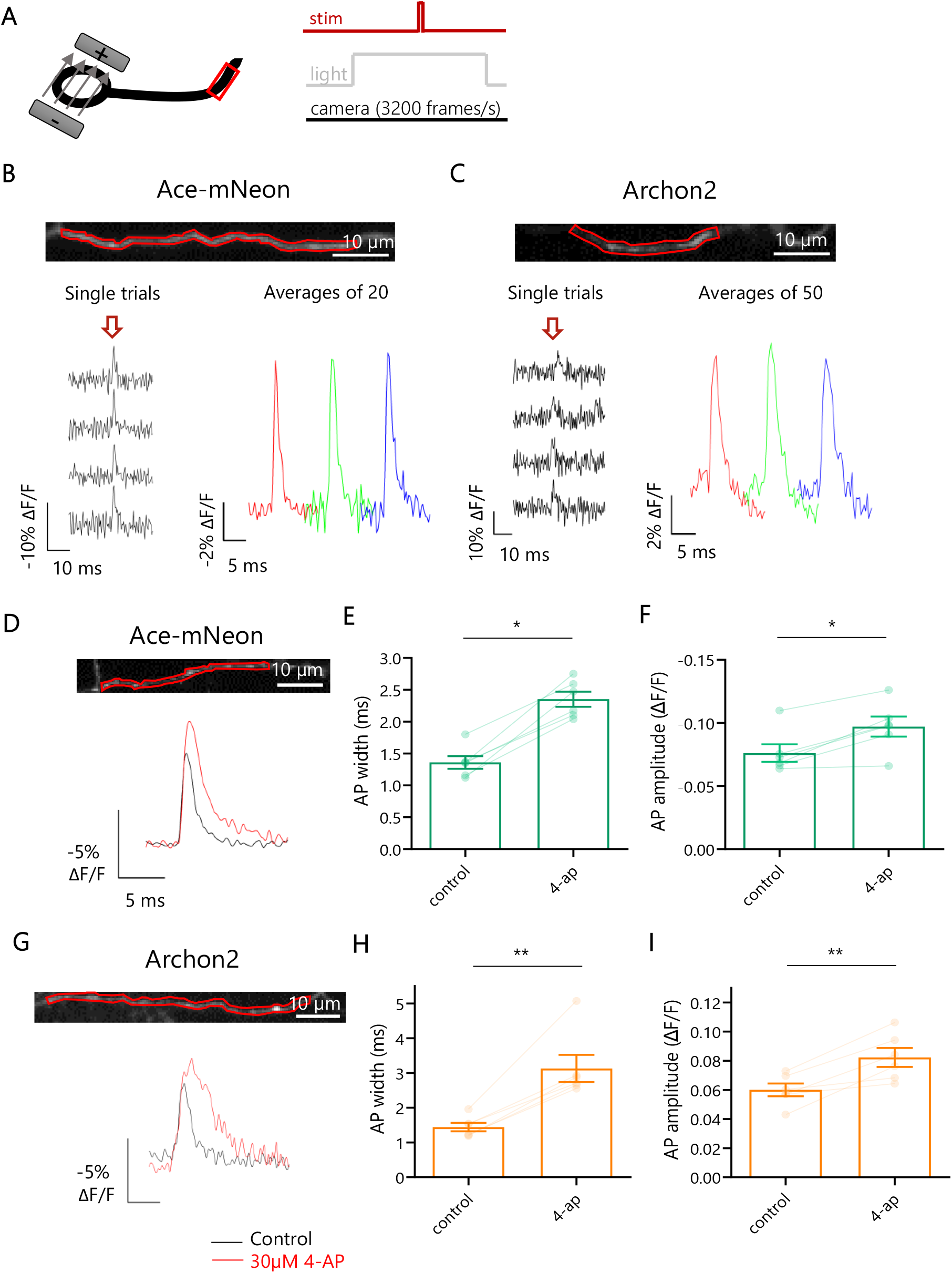
Voltage imaging of AP waveform in axons. **(A)** Schematic representation of the experimental set-up: neurons expressing Ace-mNeon or Archon2 were stimulated locally with a bipolar electrode while imaging a fragment of the axonal membrane. Time-locked 1ms stimulation pulses of 10V were delivered in order to elicit single APs, while subjecting the cells to LED or laser illumination and high-speed camera acquisition at 3.2kHz. Experiments performed at near physiological temperature (32°C). **(B-C)** Above, example axons from cells expressing Ace-mNeon (A) and Archon2 (B) with selected axonal ROIs shown in red. Below, GEVI recordings obtained from the respective ROIs, showing single trials as well as the averages over sequential groups of 20/50 repeats within the same recording. The arrow indicates the timing of stimulation pulses. **(D and G)** Above, imaging window with selected ROI of the axonal membrane shown in red. Below, average Ace-mNeon (D) and Archon2 (G) traces before and after addition of 30 µM 4-ap. **(E-F)** Axonal AP width (C) and amplitude (D) measured with Ace-mNeon before and after addition of 30 µM 4-ap. **(H-I)** Axonal AP width (H) and amplitude (I) measured with Archon2 before and after addition of 30 µM 4-ap. Ace-mNeon: N=6 cells; Archon2: N=6 cells. *, p<0.05; **, p<0.01; Wilcoxon tests.

The axonal AP waveform was similar for both GEVIs (peak ΔF/F amplitude of 6.8 ± 0.2 % and 7.5 ± 1 %, and an AP width of 1.56±0.16 ms and 1.78±0.12 ms for Ace-mNeon and Archon2, respectively; Supp. Fig.3 D and E, Mann-Whitney tests). In line with our somatic recordings (Fig 1), the axonal AP width was larger than that previously reported by electrophysiological recordings at large boutons or axons, which typically ranged from 0.3 - 1.1 ms, depending on neuron type (Geiger and Jonas, 2000; Kole, Letzkus and Stuart, 2007; Vivekananda *et al*., 2017). Such differences likely result from the on/off kinetics of the respective GEVI. As expected from Fig 1, the SNR of AP signals from Ace-mNeon was larger than that of Archon2 (27.08±3.29 vs 16.82±2.7; Supp. Fig 3 C; p<0.05; Mann-Whitney test), and was consistent with differences in fluorescence intensity between the two probes (Supp. Fig 3 F; p<0.001; Mann-Whitney test). We went on to test whether the temporal resolution of GEVI recordings in the axon was sufficient to detect AP waveform modulation. Neurons expressing either Ace-mNeon or Archon2 were stimulated locally with a bipolar electrode, and imaged before, and after perfusion with 30 µM 4-ap, to block Kv1 and Kv3 channels (Coetzee *et al*., 1999). We observed a clear widening of the axonal AP width upon application of 4-ap, in agreement with previous findings (Kole, Letzkus and Stuart, 2007; Shu *et al*., 2007; Alle, Kubota and Geiger, 2011). Notably, 4-ap also induced an increase in AP amplitude (Fig. 3 D-I; p<0.05; Wilcoxon tests). Although this observation contrasts with the electrophysiological findings in cortical neurons (Kole, Letzkus and Stuart, 2007), increases in AP amplitude have been observed in a previous voltage imaging study in hippocampal axons following blockade of the 4ap-sensitive Kv subfamilies (Hoppa *et al*., 2014).

Overall, we found that Ace-mNeon and Archon2 could reliably report modulation of the AP waveform both in the soma and in the axon. While Archon2 showed some improved kinetic accuracy over Ace-mNeon in the somatic AP waveform measurements, the superior brightness and signal-to-noise of Ace-mNeon proved better suited for axonal high-speed voltage imaging.

### Differential regulation of AP shape and Kv activity in the distal axon

Next, we performed voltage imaging experiments with Ace-mNeon in order to explore the differences in AP waveform properties in the distal axon compared to the soma and proximal axonal regions within the same cells. Neurons were subjected to local stimulation using a tungsten bipolar electrode at near physiological temperature. The evoked AP waveform was imaged at three sub-cellular locations: the somatic membrane, a proximal region of the axon (<100 µm away from the soma) and a distal region of the axon (>450 µm away from the soma) (Fig. 4 A-E). The passive propagation length constant has been found to be around 450 µm in mammalian non-myelinated glutamatergic axons (Alle and Geiger, 2006; Shu *et al*., 2006), and therefore the AP waveform measurements we made from distal axons should have experienced little to no influence from the somatic compartment. Proximal axons, on the other hand, can be subject to somatic fluctuations in membrane potential due to the close coupling between the two compartments. Importantly, proximal axons will encompass the AIS, a subcellular structure typically located between 20 and 60 µm from the soma in excitatory hippocampal cells where the AP initiates (Meeks and Mennerick, 2007; Schmidt-hieber, Jonas and Bischofberger, 2008).

**Figure 4.**
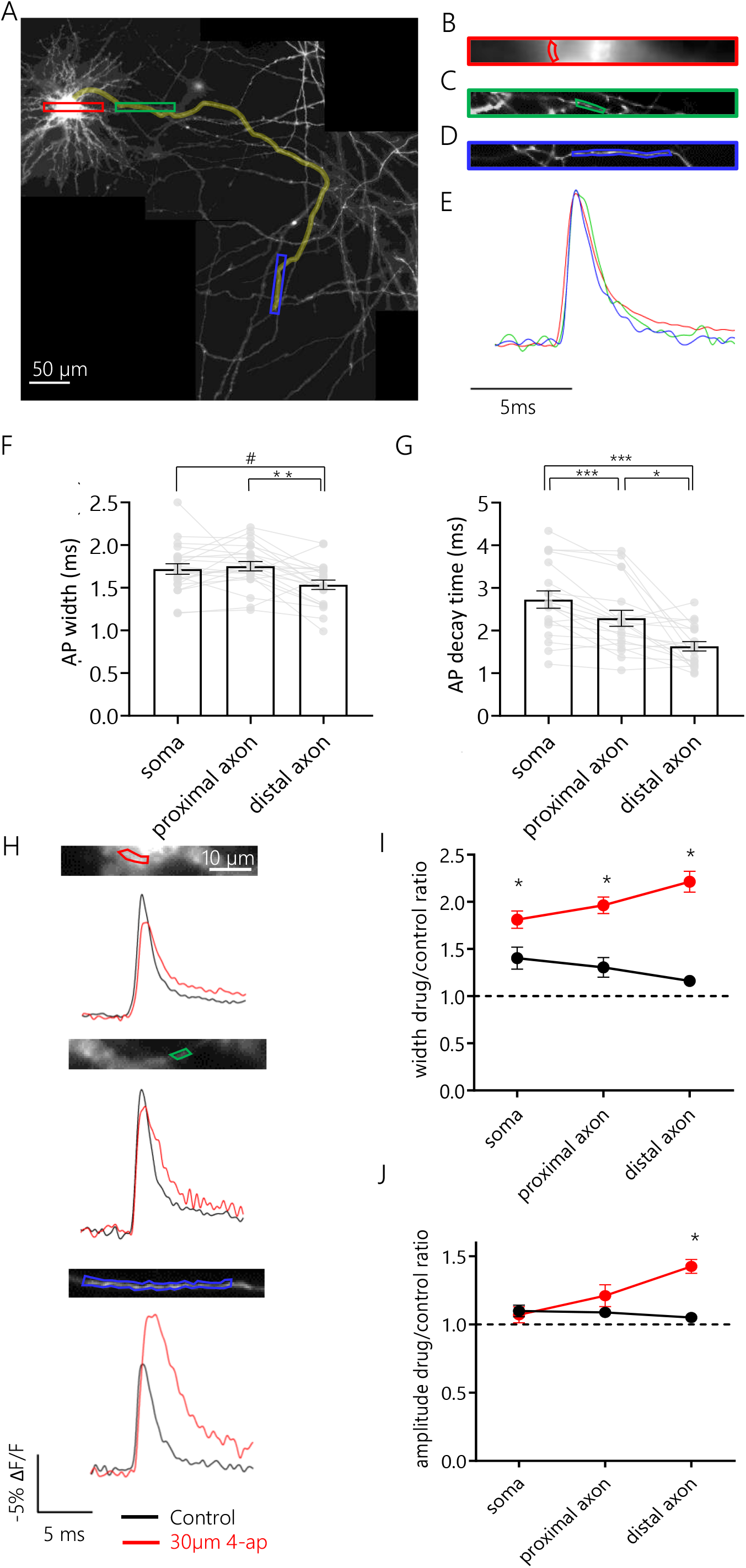
Differential regulation of AP shape by Kv channels along different sub-cellular compartments. **(A)** Reconstructed mosaic of fluorescent images of the axonal arbour of an example neuron expressing Ace-mNeon, acquired at 60x magnification. Neurons were stimulated locally with a bipolar electrode and ROIs were chosen to include a portion of the somatic membrane, a proximal and a distal segment of the axon. Highlighted in yellow, the axonal path followed up to the most distal imaged fragment of the axon. Red, green and blue rectangles indicate locations selected for imaging, enlarged in B-D. **(B-D)** Imaged sections of the somatic membrane, a region of the axon proximal to the soma, and a region of the axon distal to the soma, respectively. Images B and C are cropped from a single image series, while D was acquired separately. The ROIs drawn around membrane fragments that were selected to extract the fluorescent profile in time are shown for each image. **(F)** Aligned and overlaid averages of optical recordings of evoked APs extracted from ROIs containing the somatic membrane (red), the proximal axon (green) and the distal axon (blue) shown in B-D. **(F)** AP width recorded at the three sub-cellular locations within the same set of cells. **(G)** 80%-20% decay time of APs measured at the three sub-cellular locations within the same set of cells. **(H)** Average AP profile recorded with Ace-mNeon in the soma (above), proximal (middle) and distal (below) axon of the same cell. Traces shown for both the control condition (black) and after perfusion with 30 µM 4-ap (red). **(I)** Quantification of the effect of 4-ap (red) and mock (black) perfusion on the AP width for soma, proximal and distal axon. The width increases observed were 38% (confidence interval, CI 6% to 70%), 62% (CI 26% to 102%) and 95% (CI 085% to 148%) with respect to mock treatment, respectively. **(J)** Quantification of the effect of 4-ap (red) and mock (black) perfusion on the AP amplitude for soma, proximal and distal axon. The amplitude increase observed in the distal axon was of 32% (CI 27% to 57%) with respect to mock treatment. F and G: N=19 cells, One-way repeated measures Anova with post hoc Tukey’s multiple comparisons test. I and J: 4-ap: N=8 cells; control: N=4 cells; Mann-Whitney tests with Holm-Bonferroni correction. #, p<0.06; *, p<0.05; **, p<0.01; ***, p<0.001.

The temporal resolution of Ace-mNeon recordings (3.2 KHz) was sufficient to visualise the direction and speed of AP propagation based on latency differences between the peaks recorded at the three sub-cellular locations (Supp. Fig. 5 A and B). The speed of orthodromic axonal AP propagation was calculated from a linear fit to the latency of the distal axonal AP relative to the soma as a function of the distance from the soma, and estimated to be 383.2±53 µm/ms. This result is consistent with previous estimates of active propagation speed in hippocampal cells at physiological temperature (Meeks and Mennerick, 2007). Notably, the AP latency for proximal axonal segments was negative, indicating that AP initiation occurred in the proximal axon, presumably at the AIS.

In order to test whether the AP waveform was uniform across different sub-cellular locations, the parameters of waveforms recorded in the soma, proximally and distally within the axon were compared. Only kinetic measurements of the waveform were considered, since amplitudes cannot be reliably compared across different membrane segments without calibration (Fig. 1 E and I). Due to the rapid propagation of the axonal AP we first made sure that averaging over axonal ROIs of varying lengths was not sufficient to affect the AP shape (Supp. Fig. 5 C-F). When comparing the AP waveform across different compartments, we found that the AP was sharper in distal axons compared to the soma or proximal axons. The mean AP width decreased with distance along the axon, ranging from 1.75±0.05 ms in the proximal axon and 1.72±0.06 ms in the soma to 1.53±0.05 ms in the distal axon (Fig. 4 F). This difference was significant between proximal and distal axon waveforms and reached near statistical significance between distal axon and soma (Tukey’s multiple comparisons test after one-way repeated measures Anova, p<0.01 and p=0.056, respectively). Similar differences were observed in the 80-20% decay of the AP (Fig. 4 G, 2.73±0.2 ms, 2.28±0.2 ms and 1.63±0.1 ms for soma, proximal and distal axon, respectively, Tukey’s multiple comparisons test after one-way repeated measures Anova, p<0.001 for somatic vs. proximal and somatic vs. distal comparisons, p<0.05 for proximal vs. distal comparison). In contrast, within the distal axon, comparison of daughter and mother branches at bifurcation points did not show significant differences in width or 80-20% decay time across the different axonal branches (Supp. Fig. 6, Wilcoxon matched-pairs signed rank test).

The differences in AP kinetics observed between the somatic and axonal compartments suggested that there might be underlying differences in the channels that shape the AP in the different sub-cellular compartments. We tested the contribution of the potassium channels susceptible to block by low concentrations of 4-ap, previously shown to be targeted specifically to the axon (Kole, Letzkus and Stuart, 2007; Shu *et al*., 2007). APs were elicited by bipolar electrode stimulation in neurons expressing Ace-mNeon, and recordings were acquired at the soma, proximal and distal axon before and after bath perfusion with 30 µM 4-ap (Fig. 4 H). To control for changes in the recorded AP waveform that might occur within the experiment due to bleaching or phototoxicity, neurons from the same culture were imaged before and after perfusion with HBS without drug (mock). The results showed a strong 4-ap-induced broadening in all sub-cellular compartments, with the biggest effect on the distal axon (Fig. 4 I, p<0.05 for mock vs. 4-ap comparison in all sub-cellular compartments, Mann-Whitney tests with Holm-Bonferroni correction). Interestingly, we also observed an increase in AP amplitude with respect to the mock-only control but only in distal axons (Fig. 4 J, p<0.05, amplitude increase of 32% with respect to mock treatment, Mann-Whitney tests with Holm-Bonferroni correction). Together, our data shows that 4-ap-sensitive Kv channels play a bigger role in controlling AP shape in distal rather than proximal axons and suggests that either the activity or spatial distribution of Kv channels is biased towards distal axonal domains.

### The distal axon was resilient to frequency-dependent AP broadening

AP broadening during high frequency trains is a form of AP waveform plasticity that has been described both in the soma (Connors, Gutnick and Prince, 1982; Shao *et al*., 1999; Faber and Sah, 2003) and in large presynaptic boutons of glutamatergic neurons (Jackson, Konnerth and Augustine, 1991; Geiger and Jonas, 2000). Here, we went on to test whether the observed differences in AP properties and Kv channel activity across the axonal and somatic sub-compartments result in differential modulation of the AP waveform under stimulation trains at different frequencies. Excitatory hippocampal neurons expressing Ace-mNeon were stimulated at 20 Hz with a bipolar electrode (Fig. 5 A) and optical recordings were obtained from the soma, proximal and distal axon segments (Fig. 5 B-D). Although we observed a broadening of the AP during the train in both the soma and proximal axon (22 and 20 % increase in width of 5^th^ AP relative to 1^st^ AP, respectively), this effect was much smaller in distal axons (only a 4% increase in width) (Fig. 5E). While some significant differences in AP amplitude were observed they were generally small (less than 4% changes) and unlikely to have an important functional impact. Since the modulation of AP waveform is typically frequency dependent (Jackson, Konnerth and Augustine, 1991; Geiger and Jonas, 2000), we measured distal AP shape in response to high frequency bursts delivered at 200 Hz. At such high frequencies AP failures become much more likely (Raastad and Shepherd, 2003), so we only took recordings where we were certain that APs were fired to all stimuli. We found a broadening of the AP in distal axons at these high frequencies that was comparable to that observed in proximal axons at lower frequencies. Our results show that although distal axons were more resilient to changes in AP waveform they were capable of short-term forms of plasticity when pushed to higher frequencies. This difference in frequency tuning may also have interesting functional consequences for synaptic transmission along different axonal compartments.

**Figure 5.**
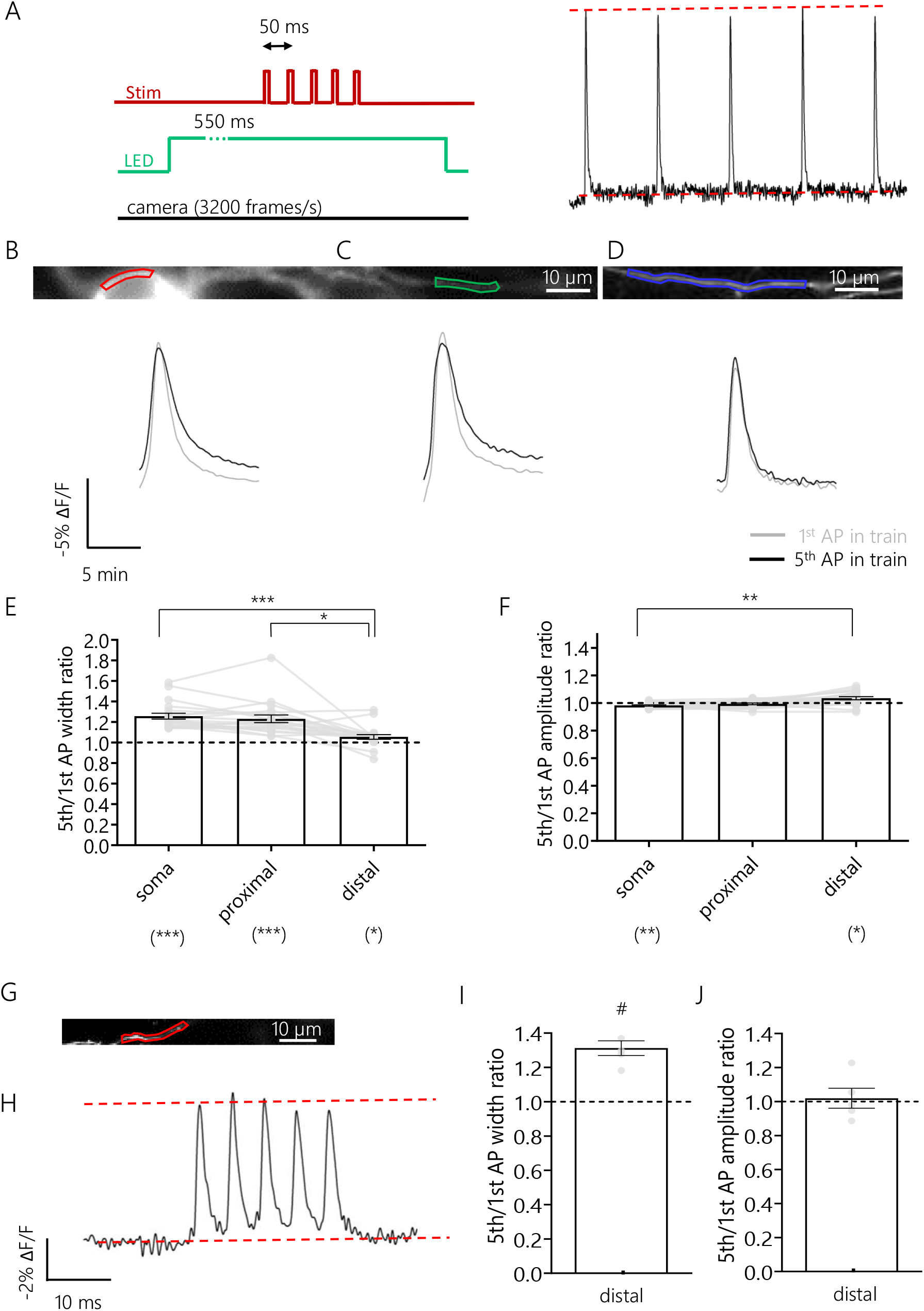
Frequency-dependent plasticity of AP waveform along different sub-cellular compartments. **(A)** Schematic representation of the experimental set-up: 5 pulse stimulation trains were delivered with an inter-stimulus interval (ISI) of 50 ms, while subjecting the cell to continuous LED illumination and camera acquisition. The 1^st^ and 5^th^ APs were compared. **(B-D)** Above, example neuron expressing Ace-mNeon with selected ROIs in the soma (B), proximal (C) and distal axon segments (D). Below, the respective average AP optic profiles of the 1^st^ and 5^th^ APs of the train overlaid. **(E)** Quantification of the width ratio between the 5^th^ and 1^st^ AP of the 20Hz train in the different subcellular compartments. The widening observed in the soma, proximal and distal axon was of 22% (CI 17.01 to 27.27%), 20% (CI 14.02 to 26.95%) and 4% (CI 2.34 to 6.98%) increase in width of the 5th AP relative to the 1st, respectively. **(F) Q**uantification of the amplitude ratio between the 5^th^ and 1^st^ AP of the 20Hz train in the different subcellular compartments. The somatic AP exhibited a decrease of 1.7% in its amplitude (CI −3.01 to −0.72%), while the AP peak in the distal axon increased by 3.8% (CI 1.0 to 6.95%). **(G)** Example axon expressing Ace-mNeon with selected ROI in red, from a neuron that was subjected to 200Hz train simulation. **(H)** Average optic trace of a 200 Hz 5-AP train recorded from the cell in A. **(I)** Quantification of the 5^th^ to 1^st^ AP width ratio. An increase of 29.7% of the 5^th^ relative to the 1^st^ AP was observed (CI 18.18 to 43.58 %). **(J)** Quantification of the 5^th^ to 1^st^ AP amplitude ratio. E-F: N=20 cells; above the graphs, Friedman test with Dunn’s post hoc multiple comparisons; below the graphs, one sample Wilcoxon tests with Holm-Bonferroni correction. I-J: N=5 cells; one sample Wilcoxon tests. *, p<0.05; **, p<0.01; ***, p<0.001; #, p=0.0625.

### Pharmacological blockade of Kv1 and Kv3 channels led to an increase in frequency-dependent AP plasticity

Our results suggest the existence of a distance-dependent difference in the short-term plasticity of AP waveform. Previous reports have implicated Kv channel inactivation in the broadening of APs during high frequency trains in axonal boutons (Jackson, Konnerth and Augustine, 1991; Geiger and Jonas, 2000). Since we showed that Kv channels played a role in controlling AP width in distal axons we next investigated whether they also played a role in controlling AP shape during a train. Imaging of Ace-mNeon was carried out in the soma and axonal domains in response to a train of APs delivered at 20 Hz, before and after perfusion of either 30 µM 4-ap or a mock control (Fig. 6 A-D). As expected, the first AP in a burst increased in width following application of 4-ap and this widening was more pronounced in distal axons. Surprisingly, however, subsequent APs broadened even further during the train, a feature that was observed along all compartments, including distal axons (Fig. 6 E, p<0.05 for all comparisons, Wilcoxon matched-pairs signed rank tests with Holm-Bonferroni correction for this and the subsequent panels). There was no change in frequency-dependent broadening in the mock perfused cells (Fig. 6 G). Finally, we observed no change in amplitude modulation upon perfusion with 4-ap or mock (Fig. 6 F and H).

**Figure 6.**
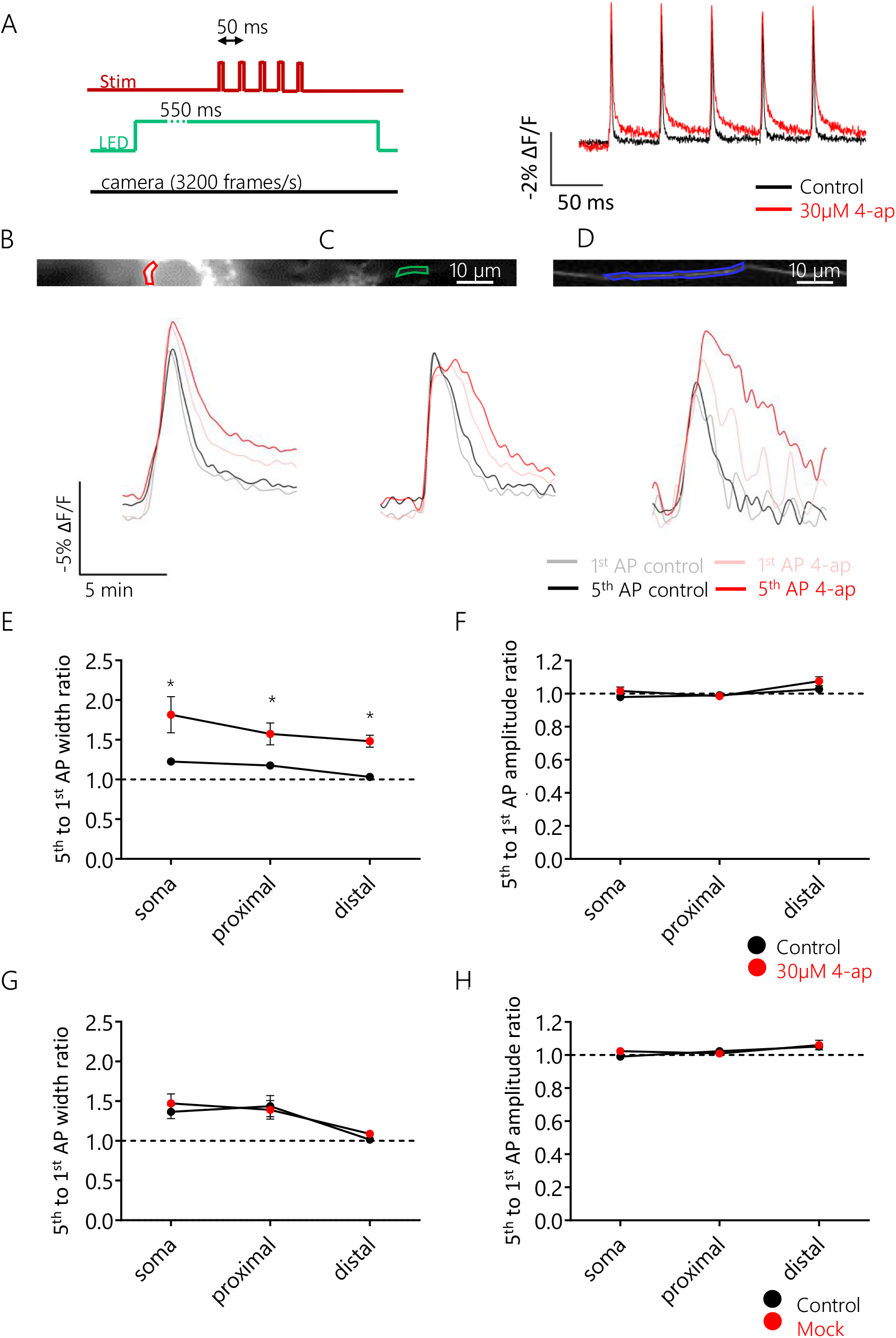
Block of 4-ap-sensitive Kv channels increased AP broadening during 20Hz trains across all sub-cellular compartments. **(A)** Schematic representation of the experimental set-up: 5 pulse stimulation trains were delivered with an ISI of 50 ms, while subjecting the cell to continuous LED illumination and camera acquisition. Cells were recorded before and after addition of 30 µM 4-ap. **(B-D)** Above, example neuron expressing Ace-mNeon with selected ROIs in the soma (B), proximal (C) and distal axon segments (D). Below, overlaid average AP optic profiles of the 1^st^ and 5^th^ APs of the train before and after addition of 30 µM 4-ap for the respective ROIs. **(E-F)** Quantification of the width (E) and amplitude (F) ratio between the 5^th^ and 1^st^ AP of the 20Hz train in the three sub-cellular compartments, before and after addition of 30 µM 4-ap. AP width facilitation increased by 30.50% relative to the control recordings in the soma (CI 10.70 to 154.9%), by 28.87% in the proximal axon (CI 2.0 to 105.0%) and by 39.51% in the distal axon (CI 11.47 to 88.15%). **(G-H)** Quantification of the width (G) and amplitude (H) ratio between the 5^th^ and 1^st^ AP of the 20Hz train in the three sub-cellular compartments, before and after perfusion with mock (HBS without 4-ap). E and F: N=8 cells; G and H: N=4 cells. *, p<0.05; Wilcoxon matched-pairs signed rank tests with Holm-Bonferroni correction.

These results suggest that 4-ap-sensitive Kv channels play a role in ensuring AP waveform stability. The increased contribution of Kv channels to shaping the AP waveform in the distal axon might therefore constitute a mechanism to guarantee faithful axonal AP propagation and reduce AP waveform plasticity.

## Discussion

In this study we have demonstrated Ace-mNeon and Archon2 were able to report the AP waveform with high fidelity, and in so doing have uncovered a role for Kv channels in controlling the shape of APs in distal axonal domains.

By performing ground truth experiments to compare somatic AP GEVI recordings with whole-cell patch-clamp recordings we showed that it was possible to capture the AP accurately and reliably using both Ace-mNeon and Archon2. Quantification of the AP waveform under control conditions, and upon drug-induced modulation of the AP shape showed that, within the same cell, changes in both AP kinetics and amplitude could be detected with GEVIs and were highly correlated to the changes observed with electrophysiology. However, when comparing across different cells only differences in AP kinetics, not amplitude, remained correlated. The most likely explanation for this is the local variation in GEVI expression, and membrane targeting, across different cells or sub-cellular compartments. Measures of AP amplitude for comparison across different cells cannot be obtained by normalising responses to resting GEVI fluorescence, as this value may not accurately represent the levels of GEVI on the membrane. Indeed, other normalising approaches are needed to obtain absolute membrane voltage measures (Hoppa *et al*., 2014). As a result, only relative changes in voltage amplitude, within the same subcellular compartment, can be compared accurately. When comparing the AP kinetics measured with Archon 2 or Ace-mNeon we found that while both sensors overestimated the AP duration, Archon2 was temporally more accurate. The most likely explanation for this difference is the faster reported on/off kinetics of Archon2 compared with Ace-mNeon (Gong et al., 2015; Piatkevich *et al*., 2018).

Compared to the soma, the ability of GEVIs to reliably report the AP waveform in the axon was much more limited by their SNR. At such low brightness levels, camera dark noise starts contributing significantly to measurement error (Popovic *et al*., 2015). In effect, GEVI AP width measurements were largely overestimated when compared with patch-clamp data from similar boutons (Geiger and Jonas, 2000; Vivekananda *et al*., 2017), even when imaged with the fast Archon2 sensor. However, both indicators were able to measure AP width and amplitude modulation upon addition of 4-ap, even in bouton-sized ROIs (∼ 1 μm^2^). Remarkably, the temporal resolution attained by voltage imaging with Ace-mNeon performed at a frame rate of ∼ 3KHz was sufficient to place AP initiation in the proximal axon and measure AP conduction velocity.

Overall, we found that the accuracy with which an AP waveform can be captured with GEVIs is affected by the filtering imposed by sensor kinetics and acquisition speed. However, the limiting factor for sub-cellular voltage imaging is SNR, which is in turn determined by sensor brightness and sensitivity (Popovic *et al*., 2015). While we found both sensors equivalent in terms of resolution and stability, there was a practical advantage in using Ace-mNeon due to its higher brightness and SNR, conducive to a higher success rate of the recordings even in cultures with variable transfection efficiency.

Taking advantage of the unique spatial resolution of GEVI imaging, we monitored the AP in three sub-cellular locations within hippocampal excitatory neurons: the somatic membrane, a proximal axon region (<100 μm from the soma) that generally encompassed the AIS, and a distal axon region (>450 µm from the soma). We observed heterogeneity in both the AP waveform and its plasticity during a train in different sub-cellular compartments, which paralleled the modulation of Kv channels in these compartments. The AP repolarisation phase was found to be shortened in the distal axon relative to the soma and proximal axon, a feature that has also been observed in other neurons including layer 5 cortical pyramidal neurons and granule cells of the dentate gyrus (Geiger and Jonas, 2000; Kole, Letzkus and Stuart, 2007). Furthermore, we found a lower susceptibility to frequency-dependent modulation in the distal axon than in the soma or the proximal axon region. Although this behaviour has been described in cortical and CA3 neurons (Meeks, Jiang and Mennerick, 2005; Kole, Letzkus and Stuart, 2007), there is evidence to the contrary in CA1 neurons, where spike broadening during a train has been shown to increase with distance from the soma (Kim, 2014). In our experiments, only high-frequency stimulation (200 Hz) resulted in the broadening of the axonal AP, a feature that may endow the distal axon with high-pass filtering properties. It has been proposed that axonal AP broadening provides a potential mechanism to modulate neurotransmission and increase the encoding capacity of the axon (Geiger and Jonas, 2000; Shu *et al*., 2006). However, the limited AP broadening in the distal axon compared to the soma observed here suggests that reliable conduction is prioritised in the axon. Maintaining a sharp AP during trains could ensure a timely membrane voltage repolarisation, minimising the inactivation of Nav channels and protecting the axon from failures (Gründemann and Clark, 2015).

The pharmacological experiments shown here revealed differential effects of 4-ap-sensitive Kv channels in shaping the AP along different sub-cellular compartments. Kv1 and Kv3 channel subtypes, susceptible to block by the low concentration of 4-ap used here, have previously been shown to control AP waveform in the axon (Kole, Letzkus and Stuart, 2007; Shu *et al*., 2007; Boudkkazi, Fronzaroli-Molinieres and Debanne, 2011; Foust *et al*., 2011; Hoppa *et al*., 2014; Kim, 2014; Rowan, Tranquil and Christie, 2014). We found that blocking Kv channels caused AP broadening across all compartments (soma and axon) but that the effect was strongest in distal axons, suggesting Kv channels may be preferentially targeted (or be preferentially activated) at these distal domains. Although all studies agree that blocking Kv channels broadens the axonal AP, the effect on AP amplitude is more controversial. A recent study using voltage imaging has shown increases in AP amplitude in the axon and implicated Kv channels in blunting the AP depolarisation (Hoppa *et al*., 2014). However, other studies, using mainly electrophysiology, have not observed any changes in AP amplitude following similar manipulations (Geiger and Jonas, 2000; Kole, Letzkus and Stuart, 2007; Alle, Kubota and Geiger, 2011). Whilst the reason for this discrepancy is not clear, and may be methodological, it is possible that differences in neuron type and position along the axon influence the outcome. Indeed, our results show changes in AP amplitude occur only in distal axons, suggesting that knowledge of axon position is a crucial parameter when assessing the role of Kv’s on AP waveform.

One surprising finding from our experiments was that the block of 4-ap sensitive Kv channels increased AP broadening during a train in all compartments measured (soma as well as proximal and distal axon). Usually, spike broadening is thought to occur through the gradual inactivation of Kv channels as the spike train progresses (Jackson, Konnerth and Augustine, 1991; Ma and Koester, 1996; Geiger and Jonas, 2000; Kim, Wei and Hoffman, 2005). Here, however, we see that following the initial increase in AP width following 4-AP application, the subsequent APs broaden further during the train. These results suggest that under basal conditions the current mediated by 4-ap sensitive channels occludes the inactivating current that would otherwise contribute to use-dependent AP broadening. In other words, Kv channels act to stabilise AP waveform from other destabilising currents. The fact that distal axons are more sensitive to 4-AP suggest that these channels are likely responsible for maintaining AP shape fidelity and invariance in these sub-cellular compartments.

The stability of the AP is particularly important in the distal axon, where up to 70% of the Nav current is inactivated (Engel and Jonas, 2005; Schmidt-Hieber and Bischofberger, 2010) and the likelihood of AP failure increases with every crossed branchpoint (Lüscher and Shiner, 1990). While a tighter control of AP kinetics might be an efficient mechanism for maintaining signal fidelity, it would be of great interest to investigate what impact it has on axonal signal processing and neurotransmitter release. A narrower AP might result in less Cav activation per AP in distal presynaptic boutons than in proximal ones. Furthermore, the differential properties of activity-dependent AP waveform plasticity in proximal and distal axon regions likely translate into further differences in neuronal output in proximal and distal boutons. Not only is the proximal axon prone to frequency-dependent AP broadening, as shown in this study, recent studies suggest that it is also susceptible to AP waveform modulation by sub-threshold signals that propagate from the soma (Rama, Zbili and Debanne, 2018). Therefore, the encoding capabilities of proximal and distal regions of the axon are likely very different. Experiments with simultaneous imaging of voltage and neurotransmitter release would reveal whether these signal processing differences translate into region-specific differences in neurotransmitter release.

## Acknowledgements

We would like to thank Thomas Knöpfel, Martin Meyer and Matthew Grubb for insightful discussions and suggestions during this project. We would also like to thank all the members of the Burrone lab for useful feedback and comments on the manuscript. Finally, we thank Kiryl Piatkevich for kindly sending us expression plasmids for both Archon 1 and 2. This work was supported by a Wellcome Trust Investigator Award (095589/Z/11/Z) and an ERC Starter Grant (282047) to JB, as well as an MRC studentship to VGS and an NC3Rs Training Fellowship to MR.

---

**Supplementary figure 1.**
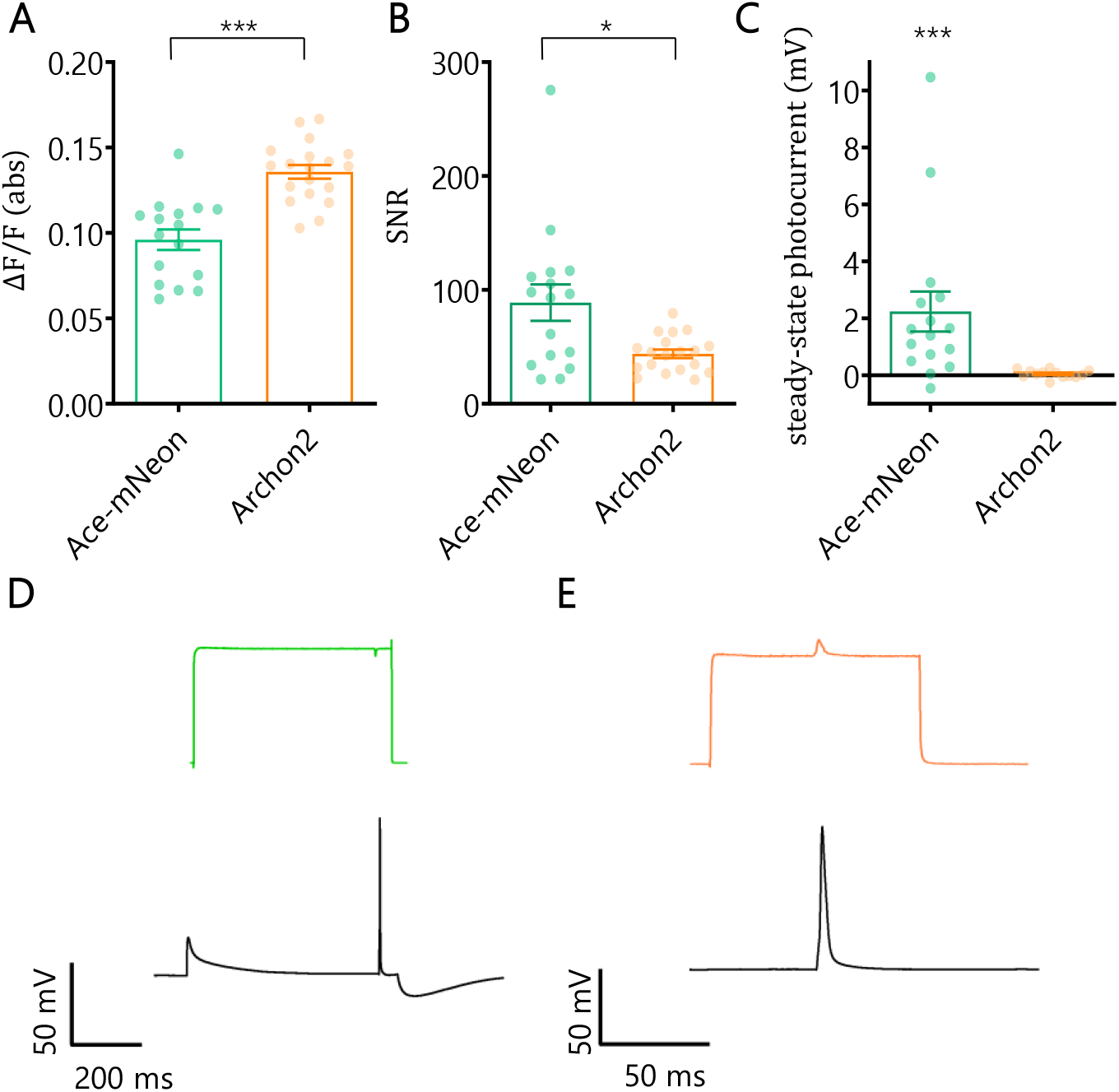
Characterisation of voltage imaging of somatic AP with Ace-mNeon and Archon2. **(A)** GEVI signal magnitude in response to an evoked AP, expressed as the absolute value of ΔF/F for Ace-mNeon and Archon2. **(B)** SNR of optically recorded APs with the two GEVIs. **(C)** Steady-state photocurrent induced in response to GEVI illumination with the corresponding light source (505 nm LED at 10 mW/mm^2^ for Ace-mNeon; 635 nm laser at 2.6 W/mm^2^ for Archon2). **(D-F)** Below, example Ephys traces acquired from neurons expressing Ace-mNeon (D) and Archon2 (E). Above, the corresponding optical recordings showing on and off LED timings to illustrate the residual photocurrent in response to light. Ace-mNeon: N=16 cells; Archon2: N=19 cells. *, p<0.05; ***, p<0.001; Kruskal-Wallis tests for A and B; one sample Wilcoxon test for C. All measurements performed on averages of 20 repeats.

**Supplementary figure 2.**
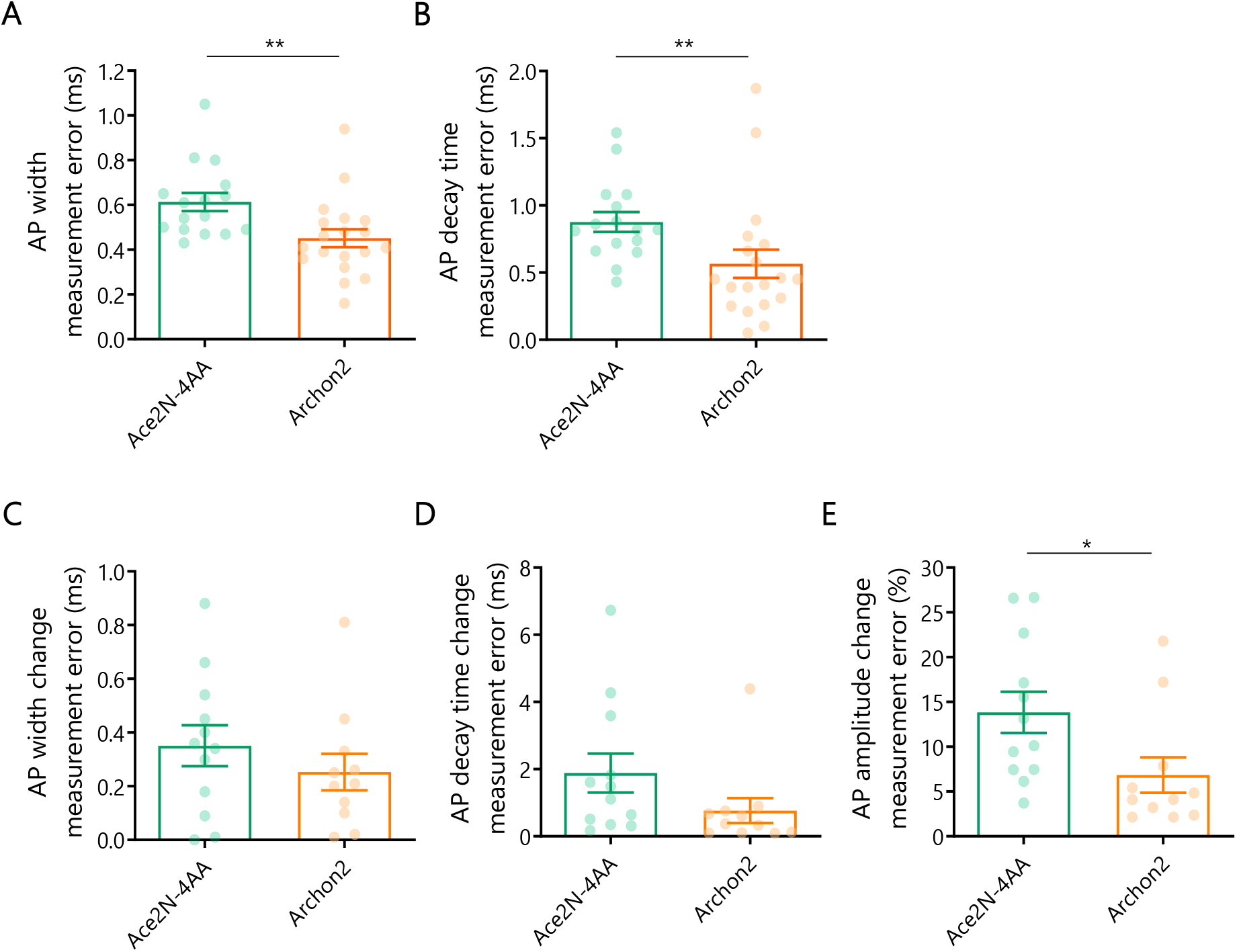
Comparison of AP parameters measurement errors for Ace-mNeon and Archon2 relative to reference electrophysiological recordings. **(A-B)** Absolute AP width measurement error (A) and decay time measurement error (B) recorded with Ace-mNeon and Archon2 relative to reference Ephys recordings. Corresponds to data in Fig. 1. Ace-mNeon: N=16; Archon2: N=19. **(C-E)** Absolute measurement error of drug-induced modulation of AP width (C), decay (D) and amplitude (E). Corresponds to data in Fig. 2. Amplitude change converted to percentage change to enable GEVI and Ephys comparison. Ace-mNeon: N=12; Archon2: N=11. All statistical comparisons performed with Mann-Whitney tests. *, p<0.05; **, p<0.01.

**Supplementary figure 3.**
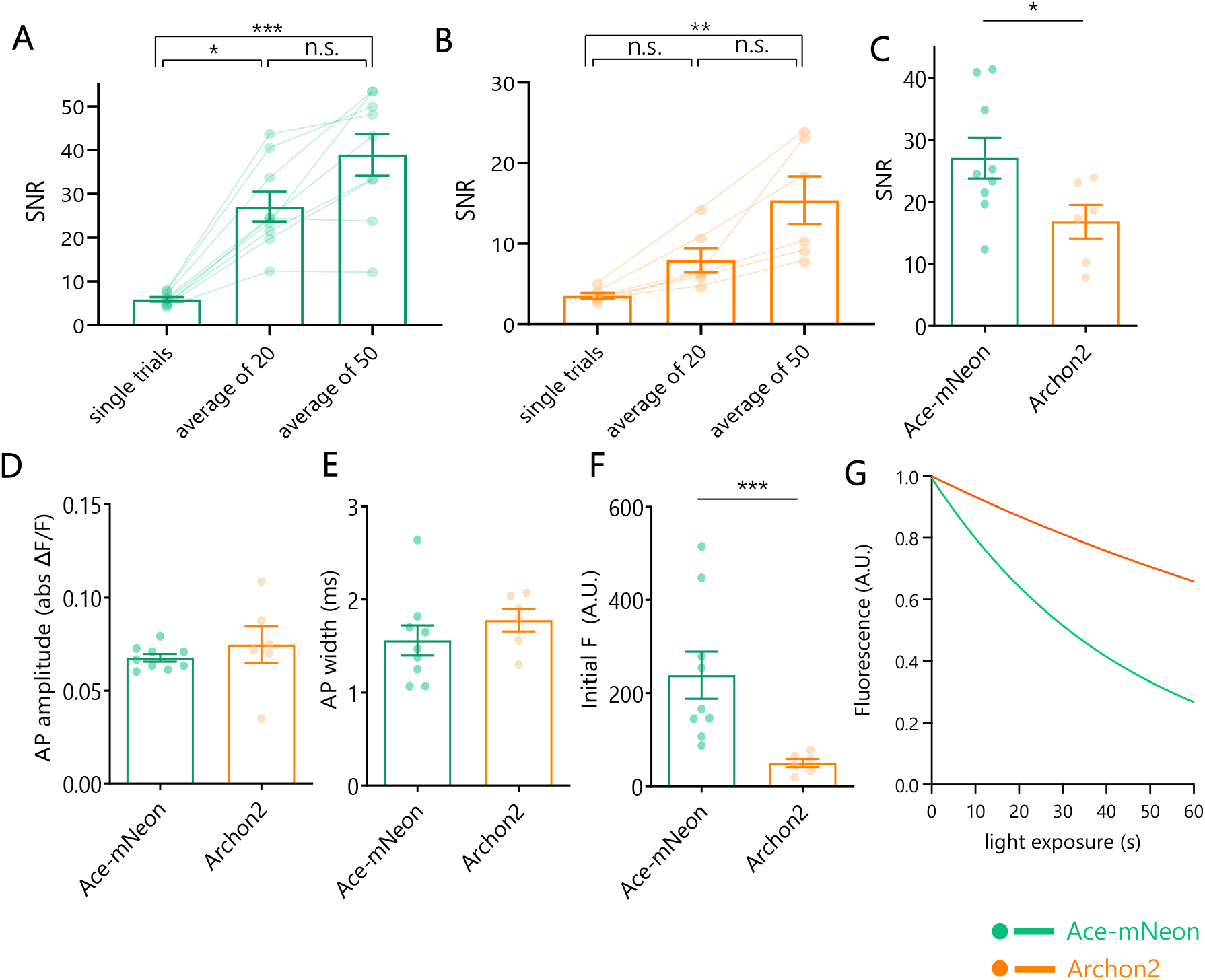
Characterisation of axonal voltage imaging. **(A-B)** GEVI AP recording SNR values for different numbers of averaged trials imaged with Ace-mNeon (A) and Archon2 (B). **(C)** SNR of axonal APs recorded with the two GEVIs. **(D)** GEVI signal magnitude in response to an evoked AP in the axon, expressed as the absolute value of ΔF/F for the two GEVIs. **(E)** Width of axonal APs recorded with the two GEVIs. **(F)** Baseline fluorescence of the first repeat under illumination with 10 mW/mm^2^ 505 LED light for Ace-mNeon and 2.6 W/mm^2^ laser light for Archon2. **(G)** Average bleaching curve of recordings with Ace-mNeon and Archon2 under exposure with 10 mW/mm^2^ 505 LED light and 2.6 W/mm^2^ laser light, respectively, normalised to initial fluorescence intensity. Ace-mNeon: N=9; Archon2: N=6; A-B: Kruskal-Wallis test with post-hoc Dunn’s multiple comparisons; C-F: Mann-Whitney tests; *, p<0.05; ***, p<0.001. Measurements in C-E performed on average recordings of 20 repeats for Ace-mNeon and 50 repeats for Archon2.

**Supplementary figure 4.**
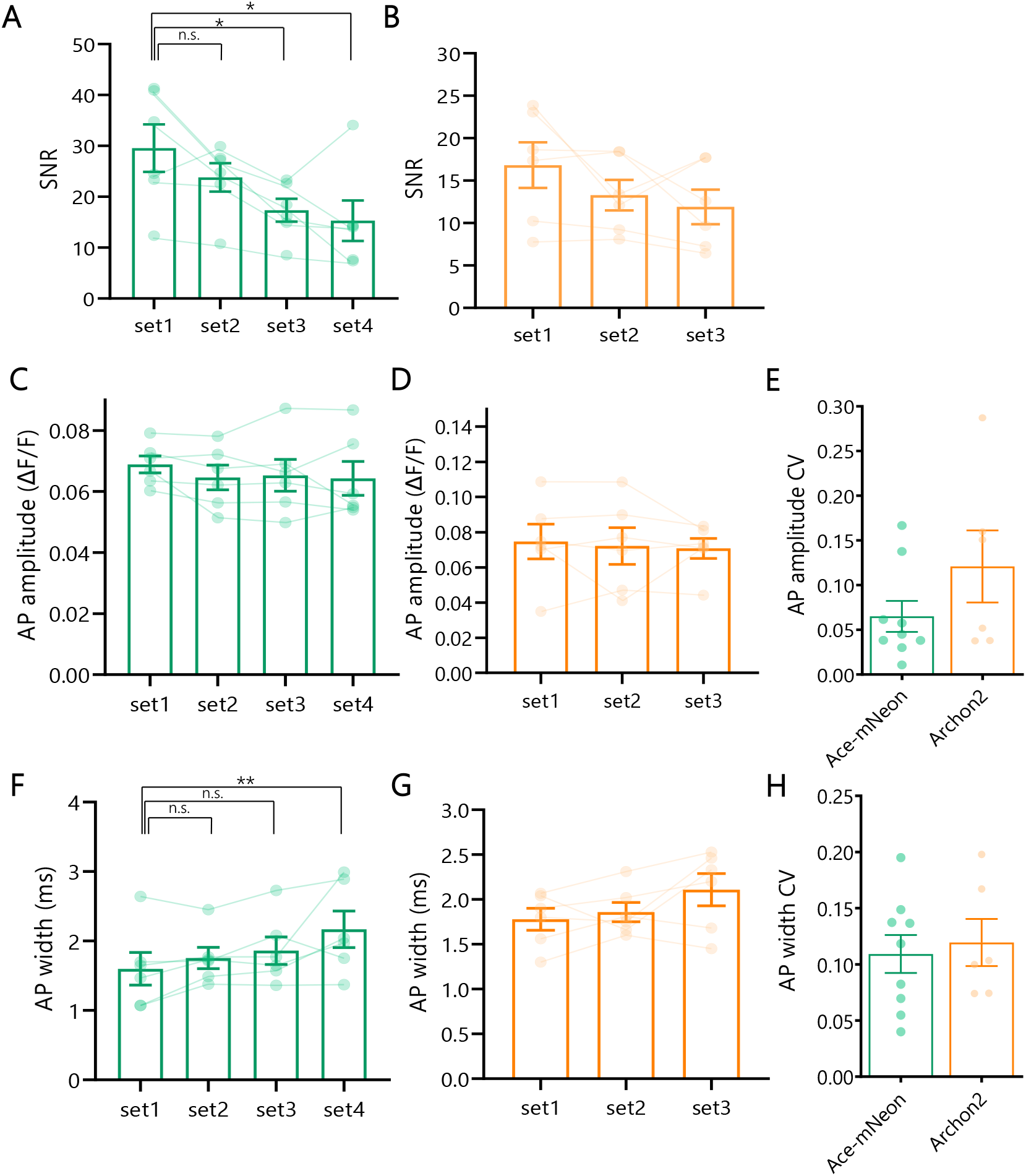
Consistency of axonal AP waveform GEVI recordings in time. **(A-B)** SNR values taken from average recordings of 4 sequential sets of 20 repeats performed with Ace-mNeon (A) and 3 sets of 50 repeats performed with Archon2 (B). Friedman tests with post-hoc Dunn’s multiple comparisons. **(C-D)** AP amplitude values measured on average recordings of sequential sets of repeats performed with Ace-mNeon (C) and Archon2 (D). Friedman tests with post-hoc Dunn’s multiple comparisons. **(E)** CV of AP amplitude values recorded in sequential sets within the same experiment performed with Ace-mNeon and Archon2, corresponding to cells shown in C and D. Mann-Whitney test. **(F-G)** AP width values measured on average recordings of sequential sets of repeats performed with Ace-mNeon (F) and Archon2 (G). Friedman tests. **(H)** CV of AP width values recorded in sequential sets within the same experiment performed with Ace-mNeon and Archon2, corresponding to cells shown in F and G. Mann-Whitney test. For A,C,F: N=6 cells; for B, D, G: N=6 cells; for E, H: Ace-mNeon: N=9 cells, Archon2: N=6 cells. *, p<0.05; **, p<0.01; n.s., not significant.

**Supplementary figure 5.**
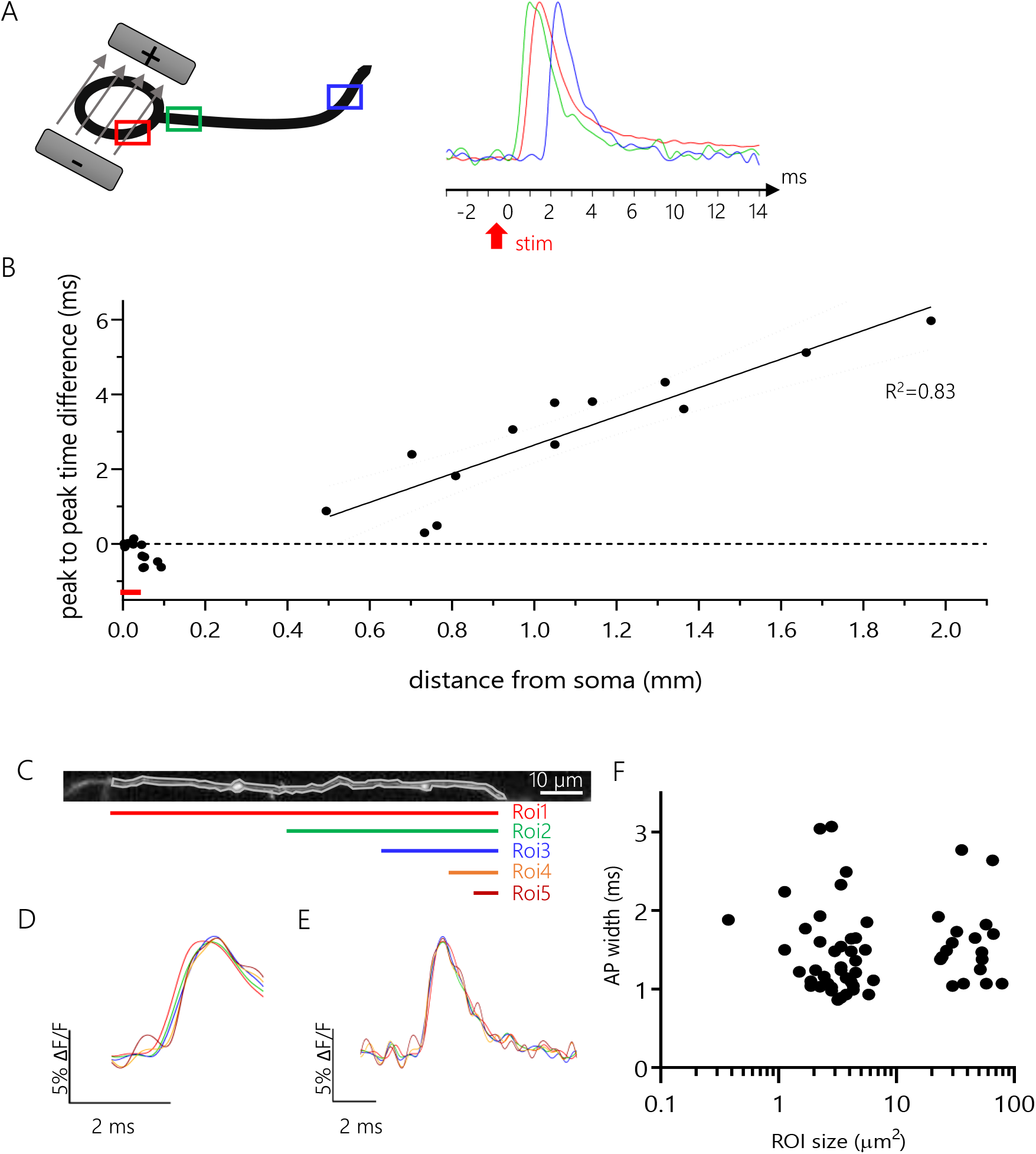
Axonal AP propagation visualised with voltage imaging. **(A)** Right, schematic representation of the experimental set up: neurons expressing Ace-mNeon were stimulated locally with a bipolar electrode and ROIs were chosen to include a portion of the somatic membrane, a proximal and a distal segment of the axon. Left, overlaid averages of optical recordings of time-locked evoked APs extracted from ROIs containing the somatic membrane (red), the proximal axon (green) and the distal axon (blue). The red arrow indicates the timepoint of stimulation with the bipolar electrode. **(B)** Time difference between the AP peaks recorded in the soma and in the axon plotted against the distance of the axon ROIs from the soma. Fit, linear regression performed with data from distal axon segments only. Red line indicates approximate location of the AIS. **(C)** Example of an imaged axonal fragment and 5 varying size ROI selections along its length. **(D)** AP rise average profiles for ROIs 1-5, illustrating delays in the rise following the direction of AP propagation. **(E)** AP average profiles for ROIs 1-5 aligned to peak do not show alteration of the AP waveform due to averaging over large sections of the axon. **(F)** Optically recorded AP width plotted against the analysed ROI size. B: N=26 axon segments (proximal and distal) from 13 cells; F: N=56 varying size ROIs from 9 cells. R^2^, goodness of fit coefficient.

**Supplemenary Figure 6.**
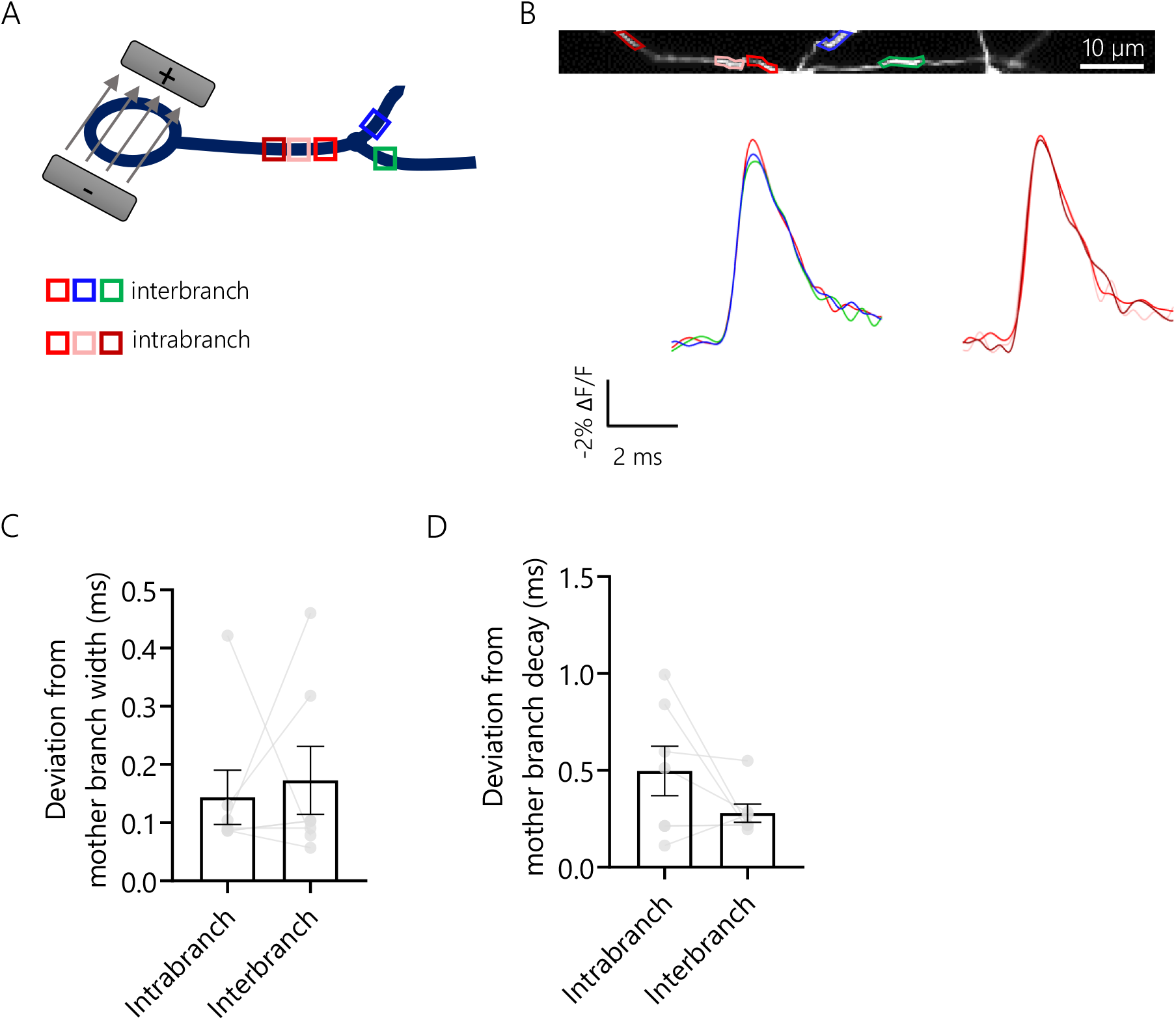
The AP waveform is not heterogeneous within the axonal arbour. **(A)** Schematic representation of the experimental set up: cells were stimulated using a bipolar electrode and an imaging window was chosen as to include an axonal branching point and assess the variability of the AP waveform obtained from ROIs selected within a single branch (intrabranch variability) and ROIs selected across different axonal branches (interbranch variability). **(B)** Example Ace-mNeon recording showing ROIs and respective average traces to visualise interbranch and interbranch variability. **(C)** Quantification of average deviation with respect to the AP width recorded at the first mother branch ROI, calculated for the other ROIs within the same branch (intrabranch) and the daughter branches (interbranch). **(D)** Quantification of average deviation with respect to the AP 80-20% decay time recorded at the first mother branch ROI, calculated within and across branches. Recordings acquired with Ace-mNeon. C and D: N=7 cells; Wilcoxon matched-pairs signed rank test.

